# Life identification number (LIN) codes for the genomic taxonomy of *Corynebacterium diphtheriae* strains

**DOI:** 10.1101/2025.06.20.660692

**Authors:** Jose F. Delgado-Blas, Martin Rethoret-Pasty, Sylvain Brisse

## Abstract

**Background:** *Corynebacterium diphtheriae*, which causes diphtheria, remains a public health concern especially in regions with low vaccination coverage. While advances in genomic typing, such as core-genome Multi-Locus Sequence Typing (cgMLST, based on 1305 genes), have improved our ability for strain identification, a standardized and stable genomic taxonomy is still lacking. This study aimed to establish a consistent classification and nomenclature for *C. diphtheriae* strains using cgMLST-based Life Identification Number (LIN) codes.

**Methods:** Comparing 1,665 genomes from *C. diphtheriae* and its closely related species *C. belfantii* and *C. rouxii*, we observed population-level genetic discontinuities in cgMLST profiles dissimilarities, and established hierarchical taxonomic levels based on optimal allelic difference thresholds. Ten-level LIN codes were defined, encompassing broad population structure subdivisions and fine-scale epidemiological levels. The LIN code system was implemented into the BIGSdb-Pasteur platform, and nicknames derived from the 7-loci MLST sequence types were given to sublineages and clonal groups.

**Results:** cgMLST genetic thresholds were first defined at species (minimum of 1,240 allelic differences) and lineage levels (1,035 differences). Sublineages (SL), clonal groups (ClG), and genetic clusters (GC) were next defined with progressively finer allelic mismatch thresholds (500, 55, and 25 differences, respectively). A broad population diversity of *C. diphtheriae* was uncovered, with the distinction of >400 SLs and >1,000 GCs. For epidemiological purposes, five shallow-level thresholds (8, 4, 2 ,1, and 0 allelic mismatches were defined, completing the 10-level LIN code taxonomy. We illustrate LIN codes applicability to investigate the genetic diversity and transmission chains of relevant clusters, such as SL8 (the 1990s ex-USSR outbreak) or SL384 (involved in outbreaks in Yemen and Europe).

**Conclusions:** The cgMLST-based LIN code system provides a stable genomic taxonomy for strains of *C. diphtheriae, C. rouxii* and *C. belfantii*. By defining ten hierarchical levels of resolution, this system effectively captures its phylogenetic diversity, facilitating population biology research and epidemiological surveillance. The public availability of this system from the BIGSdb-Pasteur platform provides a standardized framework for diphtheria genomic epidemiology with potential to harmonize global surveillance of the resurgence of diphtheria.

## Background

*Corynebacterium diphtheriae* is the classical agent of diphtheria, a severe respiratory infectious disease caused by strains that produce the diphtheria toxin (DT). DT is a potent exotoxin encoded by the *tox* gene, which is carried by a lysogenic prophage integrated into the bacterial chromosome^1^. *C. diphtheriae* strains can also cause cutaneous infections, and *C. diphtheriae* strains that are *tox*-negative can, in turn, cause invasive infections and are common in cutaneous infections. Recent work has shown that the presence of the *tox* gene in individual *C. diphtheriae* lineages can be stable during years or even decades^2–4^, highlighting the importance of precisely defining and identifying them for clinical and public health interest. Furthermore, understanding the mobility of the *tox* gene across *C. diphtheriae* sublineages, and the evolutionary relationships among *tox*-positive and *tox*-negative strains, requires a robust genome-based strain classification and naming system.

A considerable genetic diversity of *C. diphtheriae* strains was revealed by whole-genome sequencing ^2,3,5,6^. Using a core-genome MLST (cgMLST) approach, a first-generation classification and nomenclature of *C. diphtheriae* strains was proposed^3^. However, this classification has two limitations. First, it only considered two infra-specific phylogenetic levels, dubbed sublineages and genetic clusters. While useful to classify isolates into deep subdivisions and to recognize groups of very closely related isolates that may be linked by recent transmission events, this classification lacks flexibility in its definition of closely related isolates, and a single level of grouping into genetic clusters may not fit all epidemiological investigations. Second, the previous classification was based on single-linkage clustering, a mathematical algorithm that is subject to instability due to group fusion whenever intermediate genotypes are sampled. Therefore, the previously proposed classification lacks the multilevel flexibility and stability needed for a taxonomic system.

A recently developed stable strain taxonomy approach is known as Life Identification Numbers (LIN), or LIN codes^7^. LIN codes are composed of multiple hierarchical levels, which comprise numerical identifiers (integers) that capture the genetic similarity of bacterial genomes at each level. LIN codes based on cgMLST provide a proxy of genetic relationships among strains ^8,9^. Besides, groups of LIN codes can be assigned simple nicknames to facilitate human-to-human communication on bacterial strains, thus creating a strain nomenclature that bridges population biology research, public health microbiology and clinical diagnostic.

The main goal of this work was to define a cgMLST-based LIN code genomic taxonomy for strains of *C. diphtheriae* and its closely related species *C. belfantii* and *C. rouxii*^10,11^, which are often misidentified as *C. diphtheriae* in diagnostic practice. We investigated the population structure and infra-specific phylogenetic discontinuities within this species group, defined 10 hierarchical classification levels (from species level down to cgMLST identity), created a nickname nomenclature by inheritance from the 7-gene MLST identifiers, and integrated the LIN codes taxonomy into the BIGSdb-Pasteur platform. A second aim was to illustrate the benefits of the LIN codes taxonomy to address population biology and epidemiological surveillance questions in *C. diphtheriae*.

## Methods

### Database and cgMLST definition

*C. diphtheriae* is a member of the *C. diphtheriae* species complex (CdSC), which also includes, among others, *Corynebacterium belfantii* and *Corynebacterium rouxii*^10–12^. Together, these three species are referred to as *C. diphtheriae sensu lato*, as most currently used laboratory methods, including MALDI-TOF^13,14^ or PCR diagnostics^15,16^, do not distinguish them from *C. diphtheriae.* Given that all *C. belfantii* and *C. rouxii* isolates were so far reported as *tox*-negative, there is a need for genomic sequence-based identification. The BIGSdb-Pasteur platform, which relies on the Bacterial Isolate Genome Sequence Database (BIGSdb) web application and database system^17^, contains an implementation dedicated to the CdSC (https://bigsdb.pasteur.fr/diphtheria/). This resource is made of (i) genomic sequences and isolates provenance data (the isolates and genomes database), and (ii) gene sequence alleles and MLST profiles definitions (the alleles and profiles database, also known as the nomenclature database). The isolates and genomes database contains a large (> 4600 genomic sequences, January 2025) collection of high-quality CdSC genomic assemblies, their related metadata, and their genotyping information as defined in the alleles and profiles database. In turn, the alleles and profiles database hosts all the allele definitions (allele number and corresponding sequence) of the defined loci, and MLST profiles, *i.e.*, collections of loci, such as the cgMLST scheme previously defined for *C. diphtheriae sensu lato*, which comprises 1,305 loci^3^. To avoid defining cgST types with too many missing loci, each unique allelic profile is assigned to a core-genome sequence type (cgST), only when the number of missing loci does not exceed 130 loci (10%) of the total number of loci in the scheme.

For the present work, a total of 1,665 public records of *C. diphtheriae sensu lato* were used for population structure analysis: 1,611 *C. diphtheriae* genomes from previous studies^6,18^, and 44 *C. belfantii* and 10 *C. rouxii* public genomes retrieved from BIGSdb-Pasteur by April 17^th^, 2024. These records represented isolates from 50 countries located in all continents (except Antarctica), collected between 1894 and 2022, comprising a total of 1,434 distinct cgSTs.

### Population structure analyses

To assess the population structure, MSTclust v0.21b (https://gitlab.pasteur.fr/GIPhy/MSTclust) was used to generate a Minimum Spanning Tree (MST)-based clustering of cgMLST allelic profiles and to evaluate all clustering thresholds, *i.e*., from 1 to 1,305 allelic differences. All pairwise comparisons of allelic profiles between the 1,665 *C. diphtheriae sensu lato* genomes were computed using RStudio v4.3.2^19^ and the R package *tidyverse*^20^, excluding missing loci from the allelic mismatch counts. The density distribution of pairwise allelic mismatches was used to identify genetic discontinuities at different phylogenetic depths. Clustering statistics (Silhouette coefficient for the cluster consistency and adjusted Wallace coefficient for the cluster stability) were obtained for all allelic cgMLST thresholds (0-1,305 allelic mismatches) with MSTclust, and graphically visualized with RStudio using *tidyverse* and *ggplot2*^21^ packages. Allelic difference thresholds for population partitions were defined based on the distribution of Silhouette values, adjusted Wallace coefficients and allelic mismatch counts. Clustering thresholds defined by MSTclust were compared with those suggested from an analysis with pHierCC v1.24^22^.

### Creation of the LIN code taxonomy

A ten-level cgMLST-based LIN code system was established. The chosen levels were based on optimal population structure thresholds (for the fifth first ones), and shallow-level thresholds were added for epidemiological purposes (see Results). Within the LIN codes, the taxonomic resolution increases from left to right in a hierarchical manner: as the allelic distance thresholds decrease, each level is nested within the previous (left) one^7–9^.

The cgMLST-based LIN code taxonomy was implemented in the BIGSdb-Pasteur platform diphtheria database. All defined cgST genotypes (*i.e.*, corresponding to cgMLST profiles with less than 130 missing loci) were assigned a LIN code, and LIN code assignation was performed at once for the 3,247 cgST profiles (June 4^th^, 2024). The number of partitions created for each LIN code level can vary according to the profile input order, and was minimized by following an input order defined by Prim’s algorithm as implemented in the LIN codes functionality of BIGSdb^8,9^. The entire contents of the CdSC isolates database as of 26 August 2024 was incorporated into the LIN code taxonomy, representing 3,161 genomes. Circle packing charts to visualize the LIN code-based population structure were generated with RStudio v4.3.2 using the R packages *tidyverse*, *ggraph* and *igraph*^23,24^. Likewise, alluvial plots displaying the concordance of LIN code levels were generated with RStudio v4.3.2 using *tidyverse*, *ggplot2*, *ggalluvial* and *dplyr*^25,26^.

### Practical application and assessment of the LIN code taxonomy

The complete LIN codes were analyzed with the tool LINtree v0.1 (https://gitlab.pasteur.fr/GIPhy/LINtree) to generate prefix trees, which can be used as proxies to visualize the phylogenetic diversity of a dataset^9^. For comparison, actual phylogenetic trees of each dataset were constructed as follows: alignments of nucleotide sequences from all loci included in the cgMLST scheme were obtained with MAFFT v6.8^27^ applying default parameters, and then concatenated; recombinogenic positions were filtered out from the alignments using Gubbins 3.2.0^28^ with default parameters; and phylogenetic trees were generated with IQ-TREE 2 v2.3.2^29^ from the recombination-free concatenated alignments, selecting the K3Pu+F+I model with ModelFinder Plus and applying 1000 bootstrap replicates, the ultrafast bootstrap approximation and the option to optimize bootstrap support^30,31^. In addition, BactDating v1.1.2 was used to estimate the evolutionary rate and generate dated phylogenies for some specific clusters, incorporating the isolation year information from BIGSdb metadata and applying a ‘*strictgamma*’ model with 1000 MCMC iterations^32^.

LIN prefix tree-phylogenetic tree comparisons were graphically displayed as tanglegram plots performed with RStudio and using the R packages *dplyr*, *dendextend*, *ape* and *phylogram*^33–35^. Besides, one project was created for each dataset under study on the Microreact platform^36^ (https://microreact.org/) in order to integrate and visualize the LIN trees together with corresponding metadata (*e.g.*, country, isolation year, or source) and genetic traits (*e.g.*, biovar genetic determinants or *tox* allele) of the *C. diphtheriae* isolates.

## Results

### Population structure and taxonomic levels

The Silhouette consistency coefficient obtained from the 1,665 cgMLST profiles exhibited low values (∼0.1) in the 1,000 - 1,305 allelic mismatches range, which corresponded to a significant peak of pairwise allelic differences values (1,100-1,200) (Additional file 1). This peak underlines the large genomic heterogeneity of the *C. diphtheriae sensu lato* population, as a large proportion of loci had distinct alleles in pairwise comparisons. In this range, the Silhouette curve displayed two local optima, with higher consistency (Silhouette coefficient) and stability (adjusted Wallace coefficient) values, at both sides of the allelic mismatch peak, supporting the definition of two deep phylogenetic thresholds. A first peak was defined at 1,240 allelic mismatches, corresponding to the taxonomic level for species, and a second threshold was defined at 1,035 mismatches (here defined as the “lineage” level). Contrarily, in the range between 100 and 900 allelic differences, the silhouette curve reached a plateau with high consistency and stability coefficients (>0.6 and >0.8, respectively) and few pairwise allelic mismatch peaks (Additional file 1). The threshold of 500 allelic mismatches, which was previously established for sublineage (SL) definition from single-linkage clustering^3^, was located in the middle of this silhouette plateau and presented a high stability coefficient. Therefore, this threshold was retained to define sublineages (SL), with the benefit that correspondence with former SL assignations would be maximal.

Regarding the 0-100 allelic difference range, a threshold with high stability values was chosen at 55 allelic mismatches, defining a novel taxonomic level, which we defined as clonal group (ClG) (Additional file 1). The threshold at 25 allelic differences, corresponding to the previously established threshold for genetic cluster (GC) definition^3^, was supported by our current analysis (Additional file 1) and therefore retained. These five clustering thresholds were supported by pHierCC population evaluation (Additional file 2).

Last, four smaller thresholds (*i.e.*, 8, 4, 2 and 1 allelic mismatches) were selected to create high-resolution taxonomic groups that may be useful for epidemiological purposes. In all, the LIN code system provides a 10-level genome-based classification system for *C. diphtheriae sensu lato* strains (Table 1).

**Table 1.** Overview of the 10 taxonomic levels forming the cgMLST-based LIN codes for *C. diphtheriae sensu lato*.

|  | Level 1 | Level 2 | Level 3 | Level 4 | Level 5 | Level 6 | Level 7 | Level 8 | Level 9 | Level 10 |
| --- | --- | --- | --- | --- | --- | --- | --- | --- | --- | --- |
| Taxonomic assignment | Species | Lineage | Sublineage (SL) | Clonal group (ClG) | Genetic cluster (GC) | Epi level 1 | Epi level 2 | Epi level 3 | Epi level 4 | Epi level 5 |
| Minimum allelic differences<sup>a</sup> | 1240 | 1035 | 500 | 55 | 25 | 8 | 4 | 2 | 1 | 0 |
| Level left thresholds (%)<sup>b</sup> | 0 | 6.51 | 20.69 | 61.69 | 95.78 | 98.08 | 99.39 | 99.69 | 99.85 | 99.92 |
| Number of total clusters<sup>c</sup> | 3 | 30 | 388 | 834 | 1049 | 1415 | 1622 | 1835 | 1998 | 2003 |

The first five levels (blue) are based on deep phylogenetic thresholds, whereas the last five levels (green) are included for finer-scale epidemiological purposes. <sup>a</sup>Minimum allelic difference values indicate the right threshold of each level, EXCLUSIVE. <sup>b</sup>Note that 100% identical cgMLST profiles are assigned to the same complete LIN code. <sup>c</sup>Number of total clusters for each level were retrieved from the complete *C. diphtheriae sensu lato* genomic collection on the BIGSdb-Pasteur platform (August 26<sup>th</sup>, 2024).

### cgMLST-based LIN codes and nicknames for taxonomic levels

Based on the ten above-defined allelic distance thresholds (including threshold 0, which defines LIN code identity), a LIN code taxonomy was created in the BIGSdb-Pasteur diphtheria database. The population structure of *C. diphtheriae sensu lato* based on the LIN codes is illustrated in Figure 1, showing the large diversity of LIN codes and their nested relationships.

**Figure 1.**
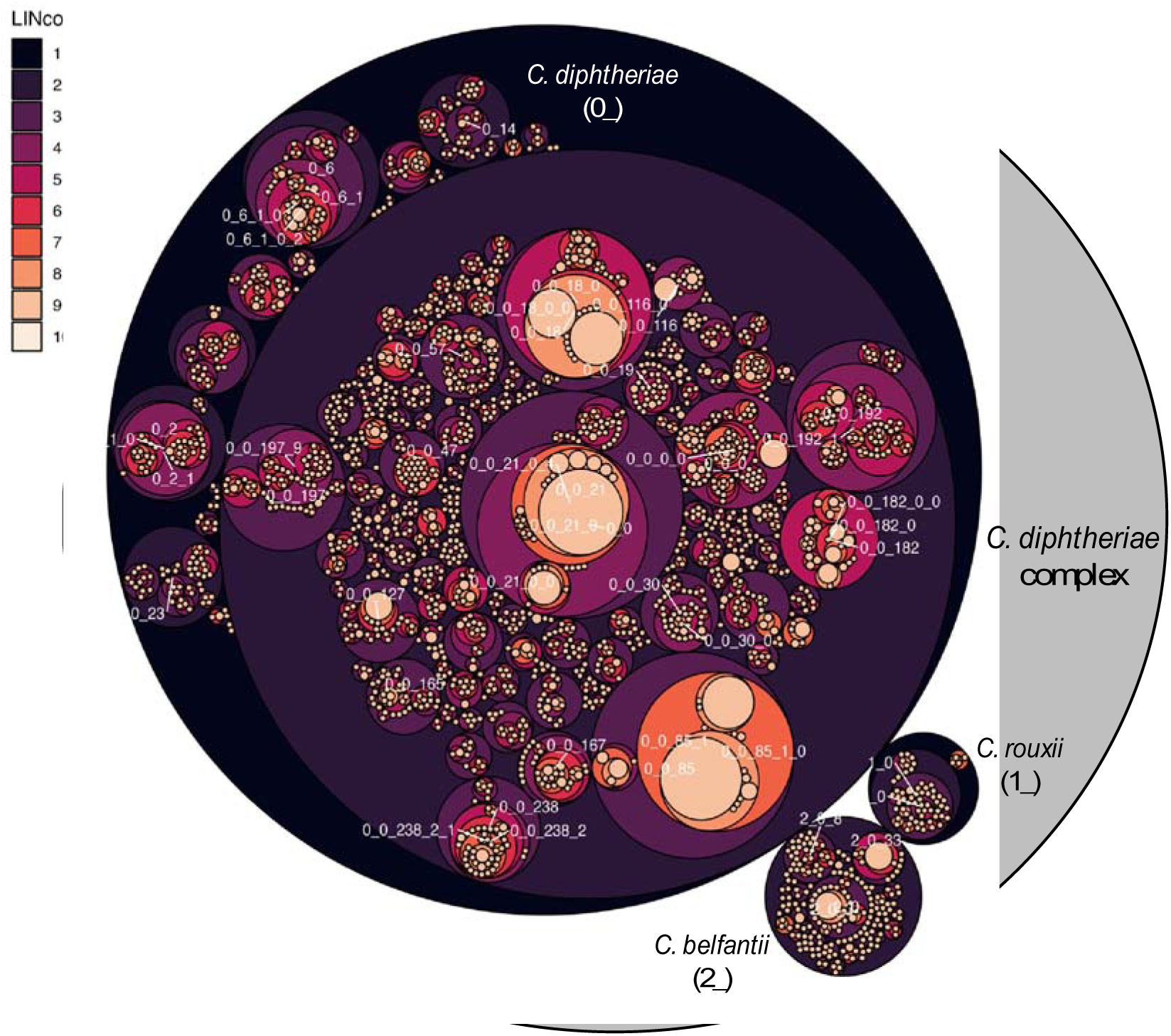
The hierarchical structure of *C. diphtheriae sensu lato* population based on LIN codes. Each bacterial strain is defined by a unique 10-level LIN code, in which each LIN code prefix is partitioned starting from 0. In this circle packing chart, each LIN code prefix is represented by a circle, which is coloured according to the level (from 1 to 10, as shown in the left legend) and nested within their corresponding higher/parental LIN code prefixes. The circle sizes display the number of genomes encompassed by each LIN code prefix, and an extra all-encompassing circle (grey) represents the entire *C. diphtheriae sensu lato* population. The first LIN code prefix corresponds to the species, as indicated in the plot, and the labels for the LIN code prefixes from level 2 to level 5 for those comprising ≥30 genomes are also displayed in the plot.

Considering the relationships between the adjacent levels of SL and ClG, whereas almost two thirds (63.5%) of the SLs were formed by a single ClG, some others comprised a larger diversity, with up to ≥10 ClGs (2.04%). When comparing LIN code levels with the STs from the classical 7-loci MLST scheme (Figure 2), a high concordance with the SL and ClG taxonomic levels was evident, confirmed by high values of the adjusted Rand indexes (*R_t_*=0.945 and *R_t_* =0.932, respectively). To facilitate communication and ensure continuity with the widely used nomenclature based on 7-gene MLST^37^, nicknames were assigned to SL (LIN code prefix of size 3) and ClG (prefixes of size 4) groups^8^ following the inheritance approach based on the most represented ST among their isolates^3^ (Figure 2).

**Figure 2.**
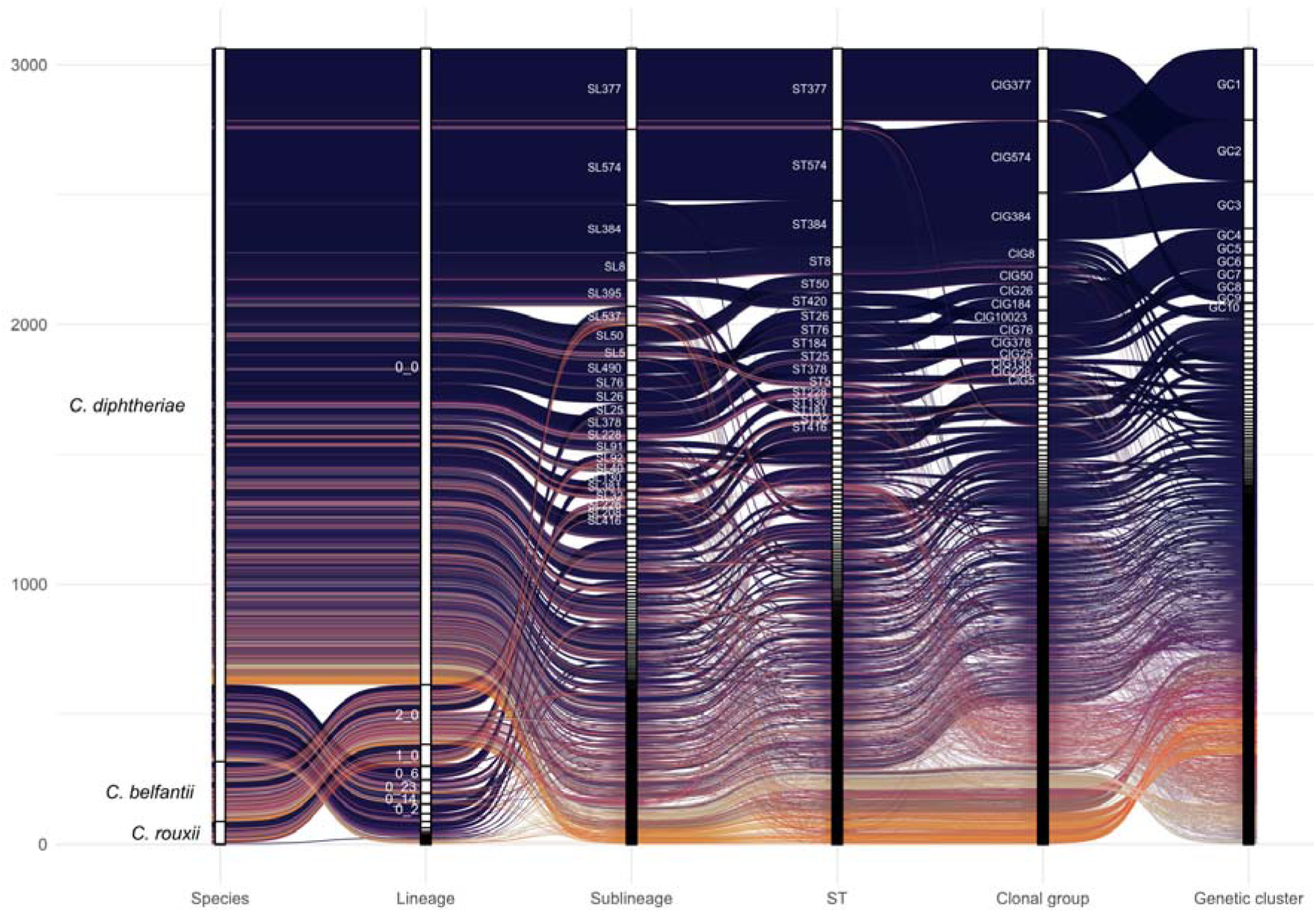
Concordance and nicknaming of *C. diphtheriae sensu lato* LIN code levels. The alluvial plot shows the hierarchical relationship of LIN code prefixes between level 1 (*i.e.*, species) and level 5 (*i.e.*, genetic cluster), and with the 7-gene MLST classification (*i.e.*, STs) (bottom labels). The inter-level links are coloured according to the different genetic clusters to facilitate the visual tracking. The nicknames for LIN code prefixes comprising ≥30 genomes are displayed next to the white bars representing each level partitions, which are ordered by decreasing total counts (*Y*-axis). Sublineage (SL) and clonal group (ClG) nicknames are based on the most represented ST among their members, as shown in the plot. The lineage level is indicated by the 2-level prefixes of LIN codes, whereas genetic cluster (GC) nicknames were assigned by decreasing total counts in the BIGSdb-Pasteur database. Alluvial plots were generated with RStudio v4.3.2 using *tidyverse*, *ggplot2*, *ggalluvial* and *dplyr*^25,26^.

In order to also facilitate communication on genetic clusters (allele difference threshold of 25; 5-level LIN code prefixes), which are useful in a broad screening approach to recognize closely related isolates potentially involved in case clusters or outbreaks^3^, GC nicknames were assigned, starting from 1, with GCs being ordered by decreasing total counts (as assessed in the BIGSdb-Pasteur database, accessed July 23^rd^, 2024). As the inheritance or total count rules can only be applied once, subsequently, novel SL, ClG and GC nicknames will be assigned by incrementing by 1 the nicknames already defined in the database. For SL and GC identifiers that were previously assigned based on single-linkage clustering^3^, the equivalence between the previously defined SLs and GCs identifiers, and the newly established LIN code-based nicknames, was made available on BIGSdb-Pasteur (see CdSC home page) and in Additional file 3.

Given that the first LIN code level corresponds to the taxonomic species level, the Linnaean taxon name of the three species was used as nickname: *C. diphtheriae* (prefix 0_), *C. rouxii* (prefix 1_) and *C. belfantii* (prefix 2_; Figures 1 and 2). We found a complete concordance between this cgMLST-based classification and the species-level ANI-derived or phylogeny-based classification of genomes.

The lineage level, represented by the 2-level prefixes of LIN codes, represented the broadest intra-species classification categories. *C. rouxii* genomes were classified into two main lineages (1_0 and 1_1) and all *C. belfantii* belonged to the same lineage (2_0; Figure 2). In turn, *C. diphtheriae* genomes were segregated into 27 different lineages (0_0 to 0_26), even though most *C. diphtheriae* (90.2%) were grouped together in lineage 0_0. Some of the *C. diphtheriae* lineages showed association with specific epidemiological traits, supporting the evolutionary significance of these lineages; for example, lineage 0_23 comprised 58 *C. diphtheriae* isolates from horses (93.6% of this lineage members), whereas no other lineage comprised isolates from this host species. In addition, lineage 0_23 was subdivided into 8 sublineages, each comprising isolates from horses, showing a concordance between their grouping at lineage level and their host source. No nickname system was created for the lineage level, as referencing them using the two-bin identifiers appears convenient.

### Main C. diphtheriae sublineages, clonal groups, genetic clusters and LIN codes

The 3-level LIN code prefixes divided *C. diphtheriae sensu lato* into 388 sublineages, including the most numerous SL377 (0_0_21, 316 isolates), SL574 (0_0_85, 300 isolates) and SL384 (0_0_18, 186 isolates), which were associated with the 2022 European outbreak in migrant populations^18^, or SL8 (0_0_0, 111 isolates), which was prevalent during the ex-USSR outbreak^3^ and is still observed sporadically nowadays (last isolate observed: year 2022).

In turn, there were a total of 834 clonal groups (Table 1). A few clonal groups belonging to the aforementioned sublineages were overrepresented in the database, accounting for more than a quarter of the total records: ClG377 (0_0_21_0), ClG574 (0_0_85_1), ClG384 (0_0_18_0), and ClG8 (0_0_0_0) (Figures 1 and 2).

The LIN code taxonomy further distinguished a total of 1,049 genetic clusters, which correspond to the maximum level of genetic variation previously observed among documented *C. diphtheriae* outbreaks^3^. As expected, some frequent genetic clusters were observed within the most frequent sublineages and clonal groups, and the three main ones belonged to ClG574 (GC1, 0_0_85_1_0), ClG377 (GC2, 0_0_21_0_4) and ClG384 (GC3, 0_0_18_0_0; Figures 1 and 2)^18^. Contrarily, the diverse ClG8 was split up in multiple GCs, reflecting the longer evolutionary history of this clonal group.

The epidemiological levels defined for surveillance purposes further divided the genetic clusters into finer groups, from the 6-level prefixes (1,415 groups) to the complete 10-level LIN codes (2,003 groups) (Figures 1 and 2; Table 1). We illustrate below the use of LIN codes to investigate genotype-phenotype links and potential transmission within specific genetic clusters and epidemiological contexts.

### Biovar and tox gene associations with C. diphtheriae taxonomic levels

Based on phenotypic properties (nitrogen reduction and glycogen or starch utilization), the classification of *C. diphtheriae* strains into biovars has been performed since the 1930s. Recently, these distinct biochemical profiles were linked to specific genetic markers: gene *spuA* and the *narIJHGK* gene cluster^6^. We evaluated the distribution of these genetic markers in the *C. diphtheriae* population using the LIN code classification of a dataset of 1,250 *C. diphtheriae* isolates previously characterized for their biovar and the two genetic markers^6^. The *spuA* gene was strongly associated with all LIN codes levels, starting from the *C. diphtheriae* lineage level. All lineages but one, 0_0, were uniform for this marker status (96.0%). At the sublineage level, the proportion of *spuA*-uniform groups was 97.9%; only five out of 202 sublineages from lineage 0_0 (*i.e.*, SL5, SL122, SL228, SL295, and SL466) possessed both *spuA* profiles. The majority of 0_0 sublineages therefore only comprised either *spuA*-positive (33.15%) or *spuA*-negative (66.85%) isolates. In turn, the *spuA* profile was almost fully concordant with the clonal group level, except for one clonal group (ClG466) out of the total 464 (99.7%), which was formed by a main *spuA*-positive genetic cluster and two one-isolate *spuA*-negative genetic clusters. Therefore, LIN code classification at clonal group level is concordant with *spuA* presence with only one exception, and thus appears useful to predict biovar (Gravis, *spuA*-positive; *versus* Mitis or Belfanti, molecularly defined as *spuA*-negative). Last, the distribution of *spuA* was fully concordant with genetic clusters (583/583, 100%), thus implying a complete biovar homogeneity at the evolutionary scale of outbreaks or epidemiological clusters.

Similarly, when looking at the *tox* gene presence and allele number as a function of LIN codes, we found a strong homogeneity at the level of sublineages (*tox* presence: 89.4% of homogeneous groups; allele: 84.6%), clonal groups (*tox*: 97.6%; allele: 95.9%) and genetic clusters (*tox*: 98.6%; allele: 97.9%). This demonstrates the utility of these taxonomic levels as predictors of the *tox* status and even of the *tox* allele (Figure 3, Additional file 4). In other words, most *tox* alleles were strongly associated with specific sublineages or clonal groups. Nevertheless, some Mitis sublineages exhibited a broad diversity of *tox* alleles, particularly SL141 (*tox5*, *tox13* and *tox22*), SL381 (*tox2*, *tox3*, *tox18* and *tox20*) and SL40 (*tox2*, *tox3*, *tox27* and *tox37*) (Additional file 4). Four *tox* alleles were identified in more than five distinct sublineages (*i.e.*, *tox2*, *tox3*, *tox8* and *tox16*), including the most prevalent one, *tox2*, present in 112 isolates spanning 26 different SLs (Figure 3), suggesting a successful dissemination through horizontal gene transfer.

**Figure 3.**
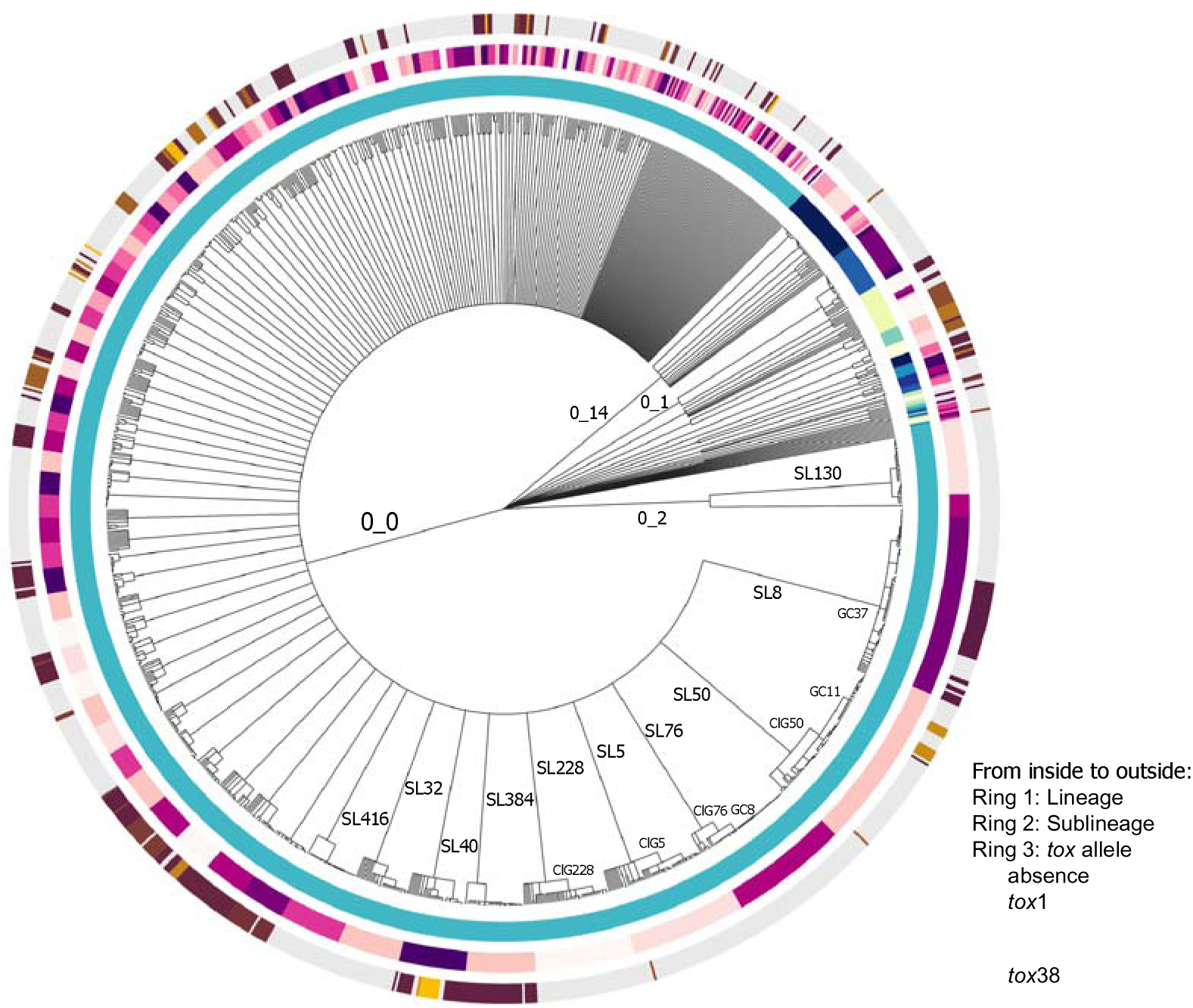
Distribution of taxonomic clusters and *tox* alleles in the *C. diphtheriae* population. The LIN tree in the center was generated using cgMLST-based LIN codes from a collection of 1,250 genomes previously published^6^. Concentric rings show taxonomic clusters and *tox* allele presence according to the legend on the right. Different lineages and sublineages are represented by alternate colours, and relevant lineages, sublineages (SL), clonal groups (ClG) and genetic clusters (GC) are indicated by labels next to their corresponding tree branches.

Several recent studies have identified a higher *tox* gene prevalence in biovar Mitis isolates than in biovar Gravis ones (*or spuA*-positive isolates)^6^. Here, the proportion of *tox*-positive isolates was similar for both *spuA*-positive (28.5%) and *spuA*-negative (31.7%) isolates, but the *spuA*-negative isolates displayed a much broader diversity of *tox* alleles (32 allelic variants) compared to the *spuA*-positive ones (6 allelic variants). Some narrow-range alleles were only found in all members of specific Gravis lineages (*e.g.*, *tox12* in 0_6 or *tox21* in 0_11), but unlike at shallower LIN code levels, the *tox* status and allele predictions according to the lineage were limited (76.0% and 64.0%, respectively) (Figure 3).

### Application of LIN codes to epidemiological studies of C. diphtheriae

We first explored the LIN codes diversity of SL8, based on 91 isolates, which were mainly collected in Europe (Figure 4A). SL8 includes ex-USSR outbreak isolates^3^. Here, SL8 comprised a unique clonal group (ClG8) and all isolates were of 7-gene MLST type ST8, but there were 15 genetic clusters (ten of which comprised less than 10 isolates), highlighting the genetic diversity of SL8. The LIN codes prefix tree displayed a high concordance with the SNP-based phylogenetic tree (Additional file 5), thus providing a proxy to visualize isolates relationships. The *tox* gene status within SL8 was fully concordant with the genetic cluster segregation, with *tox1* being the only *tox* allele in *tox*-positive GCs (Figure 4A). Of the 15 GCs within SL8, some were highly clonal, such as GC32, in which all isolates shared the same 7-level LIN code (0_0_0_0_5_0_0) and were collected between 2016 and 2017 from several German regions, suggesting clonal dissemination (Figure 4A and 4B). On the other hand, other genetic clusters were more diverse, such as GC37 (0_0_0_0_2), whose members were isolated in Belarus over a period of 11 eleven years, or GC29 (0_0_0_0_0), which included isolates obtained in five different European countries, revealing international spread (Figure 4A and 4B). Within some countries, SL8 isolates were genetically diverse, implying several transmission chains: 43 isolates from Germany belonged to five GCs, and from Belarus, eight distinct GCs were identified (Figure 4B). This genetic diversity likely reflects the persistence of SL8 and its diversification since the 1990’s ex-USSR outbreak; for example in Belarus, SL8 was isolated at least from 1996 to 2010^38^.

**Figure 4.**
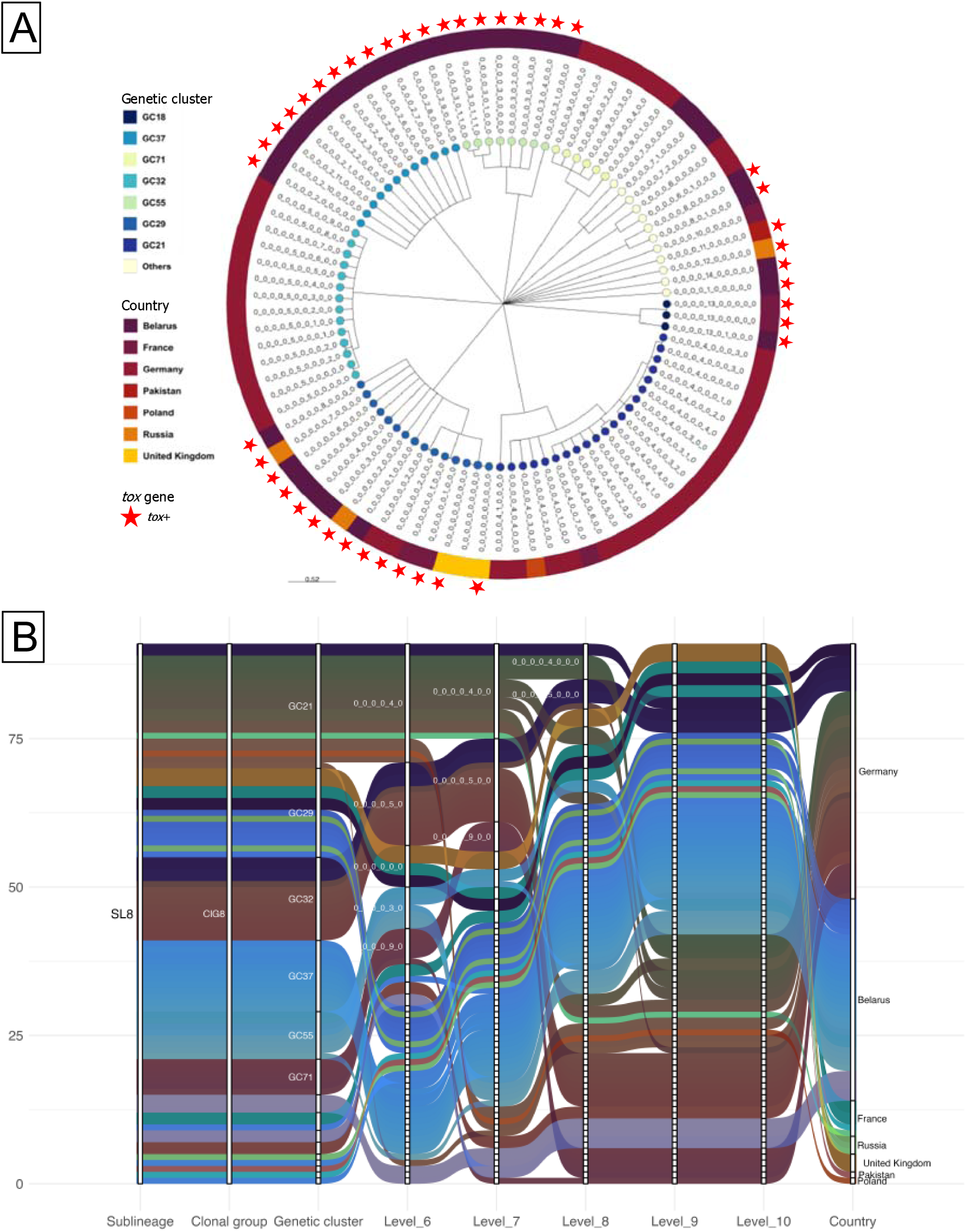
LIN codes in the highly diverse *C. diphtheriae* SL8. **A.** LIN tree of the 91 included genomes. Tree tips are coloured according to the distinct genetic clusters (GC) (left legend), and leaf labels show the complete 10-level LIN codes. The country of isolation is indicated by the different ring colours (left legend), and the *tox* gene presence is indicated by a red star. **B.** Alluvial plot displaying the partition relationships based in LIN code prefixes from level 3 (*i.e.*, sublineage) to level 10 (*i.e.*, complete LIN code) (bottom labels), and the country of isolation (right labels). The inter-level links are coloured according to the different complete LIN codes and countries. The nicknames for LIN code prefixes comprising ≥5 genomes are displayed next to the white bars representing each level partitions, which are ordered by decreasing total counts (*Y*-axis).

We next revisited the endemicity and local transmission of *C. diphtheriae* in New Caledonia between 2015 and 2019, based on genomic data from a previous genomic epidemiological study^39^. The 58 *C. diphtheriae* isolates belonged to four distinct lineages, revealing the deep population diversity existing in the New Caledonia archipelago. As in the total dataset, lineage 0_0 was the most prevalent in New Caledonia, with 43 members, and was highly diverse, with 13 sublineages and 22 genetic clusters (Additional file 6A). The LIN codes prefix tree showed a high concordance with the phylogenetic structure of the population (Additional file 6A); however, there were exceptions where distinct LIN code lineages were located in the same deep branch of the cgSNP tree, such as 0_11 and 0_0. The SLs with the highest isolates counts were SL228 (0_0_30, n=21) and SL416 (0_0_47, n=16), which encompassed 10 and 1 GCs, respectively, illustrating how LIN codes are concordant with their different epidemiological patterns described in the original work: SL228 presented a wider geographical distribution compared to SL416.

We next investigated LIN codes in SL384, an epidemiologically important *C. diphtheriae* sublineage, which was predominant in the Yemen outbreak and also frequently isolated during the 2022 outbreak in asylum seekers in Europe^18^. A total of 183 SL384 genomes were included, all collected in European countries (2022-2024) and in Yemen in 2018^40^. All SL384 members carried *tox2*. SL384 was subdivided into three clonal groups and their corresponding three genetic clusters. However, two of these were rare, comprising only one (GC530) or two (GC208) isolates. In contrast, GC3 comprised all other SL384 isolates (Figure 5A and 5B), suggesting a recent clonal expansion. Furthermore, GC3 included two main complete LIN codes that differed by only two allelic differences (0_0_18_0_0_7_0_0_0_0 and 0_0_18_0_0_7_0_0_6_0), which comprised most (93.4%) of the isolates belonging to the European outbreak, consistent with a very recent common ancestor, estimated in 2019 by Hoefer *et al.*^18^ In contrast, the isolates from Yemen were distributed in 15 different 7-level LIN code prefixes (4 allelic differences), showing the broader GC3 diversity in Yemen (Figure 5A and 5B). The contrasted diversity of LIN codes between Yemen and European outbreaks suggests two distinct epidemiological scenarios, namely an endemism of GC3 in Yemen versus a rapid spread in Europe. This case illustrates how LIN codes can be used to infer distinct epidemiological dynamics.

**Figure 5.**
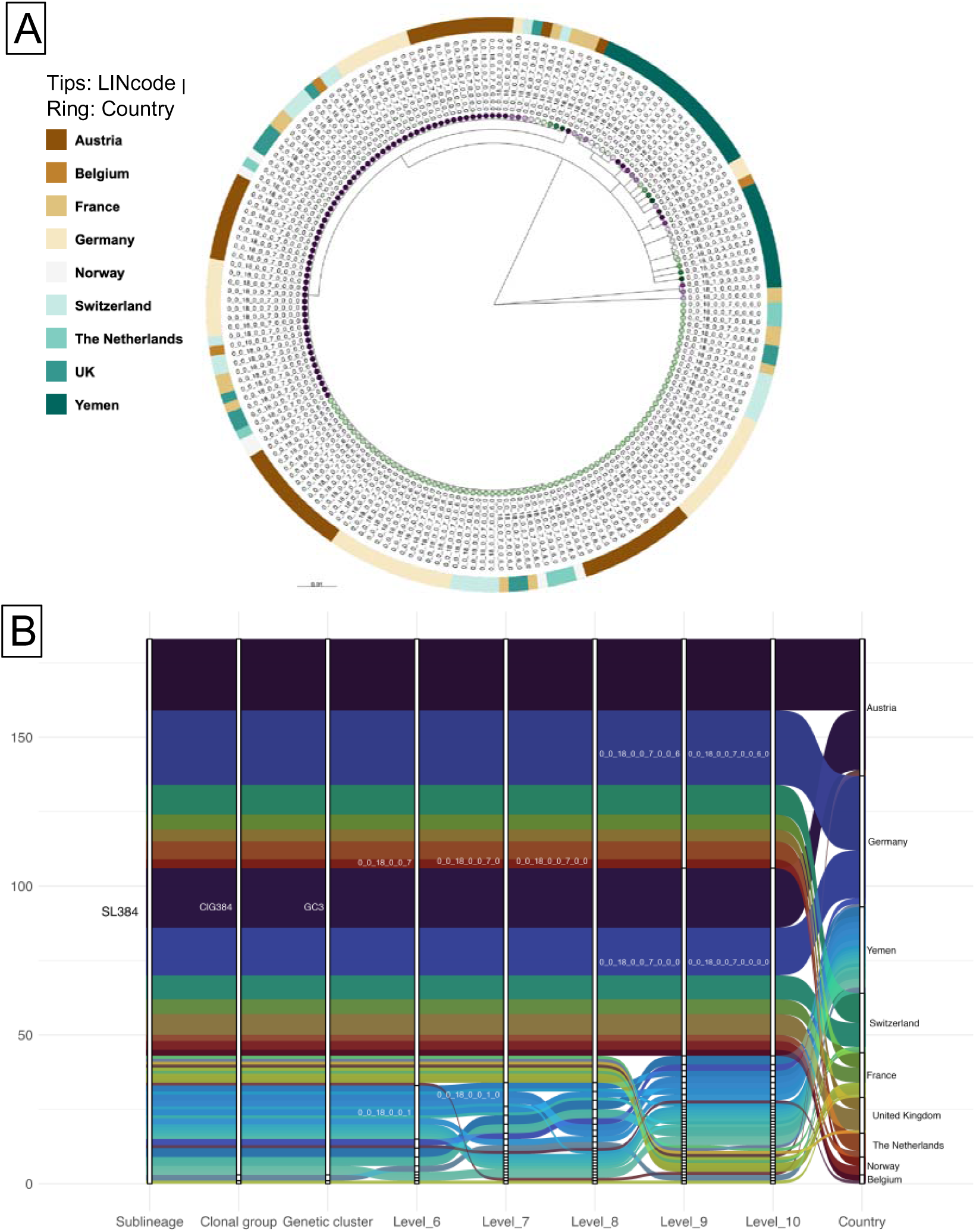
LIN codes in the highly conserved *C. diphtheriae* SL384. **A.** LIN tree from a collection of 183 genomes from CdSC BIGSdb-Pasteur. Tree tips are coloured according to the distinct 9-level LIN code prefixes, and tree labels show the complete 10-level complete LIN codes. The country of isolation is indicated by the different ring colours (left legend). **B.** Alluvial plot displaying the cluster distributions based in LIN code prefixes from level 3 (*i.e.*, sublineage) to level 10 (*i.e.*, complete LIN code) (bottom labels), and the country of isolation (right labels). The inter-level links are coloured according to the different complete LIN codes and countries. The nicknames for LIN code prefixes comprising ≥5 genomes are displayed next to the white bars representing each level partitions, which are ordered by decreasing total counts (*Y*-axis).

As a fourth example, we looked into the LIN codes diversity of GC1, which was also one of the main clusters responsible for the 2022 European outbreak^18^. GC1 (0_0_85_1_0) belongs to SL574 (0_0_85), which was so conserved that GC1 members represented almost all isolates of SL574 (94.6%), distantly followed by GC26 (4.8%) and GC537 (0.3%). From the 140 GC1 genomes available, 10 distinct complete LIN codes were identified, six of which only comprised a single isolate (Additional file 7A). Notably, Switzerland and France were the reporting countries of four and three of these single-isolate LIN codes, respectively, showing higher GC1 diversity than other countries (Additional file 7B). Still, only two LIN codes comprised the vast majority of GC1 members: the main one was 0_0_85_1_0_0_0_0_0_0, which encompassed 92 isolates (65.7% of GC1) and was present in all participating European countries, except Italy; whereas 0_0_85_1_0_0_0_2_0_0 comprised 37 isolates (26.4%) and was distributed in all countries except for Norway (Additional file 7B). The observation of identical LIN codes in multiple countries illustrates the efficiency of this system to track the rapid spatial spread of outbreak-causing clones.

Last, to investigate how *C. diphtheriae* LIN codes change during recorded transmission chains, we revisited in light of LIN codes, a long-term local transmission of an NTTB strain in England^4^. Eight isolates were collected between 2014 and 2021, in a rehabilitation skin clinic and households of patient(s). The eight isolates belonged to ST336, but nevertheless showed considerable phylogenetic diversity, with 7-199 SNPs and 3-109 cgMLST mismatches^4^. LIN codes show they belong to five distinct genetic clusters, two of which were detected more than once: GC1071 (0_0_246_0_0), with three isolates collected from the same patient, and GC1070 (0_0_246_0_1), which comprises two isolates from two household contacts of the mentioned patient (Additional file 8A). Three genetic clusters belong to the same clonal group (ClG10037, 0_0_246_0), and have originated from a most recent common ancestor (MRCA) estimated in the late 1990’s (Additional file 8B), which in turn belong to SL10037 (0_0_246). This sublineage encompasses three clonal groups involved in this epidemiological investigation (Additional file 8A), and exhibits an evolutionary rate of 1.3x10_-6_ substitutions/site/year (95%CI 5.4x10^-7^-2.4x10^-6^). It was estimated to have originated in the mid 1960’s (MRCA: 1964, 95% CI 1942-1991) (Additional file 8B), consistent with the large genomic diversity observed among the isolates.

## Discussion

Taxonomic classification and nomenclature are needed for communication on bacterial strains in research and epidemiological surveillance. *C. diphtheriae* comes back as a significant public health threat, particularly in low- and middle-income countries (LMICs) with suboptimal vaccination coverage, and antimicrobial-resistant strains are emerging^18,41^. Previous work has focused on identifying *C. diphtheriae* phenotypic and genotypic groups, including through biotyping and other pre-molecular approaches. The 7-loci MLST scheme provided initial genotypic classification, but lacks the resolution needed for fine discrimination^3^. The 1,305-loci cgMLST extension of this approach now enables high-resolution characterization of *C. diphtheriae* isolates. While standardized classification and naming at multiple levels is needed for effective use of cgMLST data, such a taxonomic system has been lacking. Here, we developed a 10-level LIN code taxonomy for *C. diphtheriae, C. belfantii* and *C. rouxii* strains, and explored how the LIN code approach based on cgMLST data could help addressing epidemiological and evolutionary questions, as shown previously for other relevant bacterial pathogens, including *Klebsiella pneumoniae* and *Streptococcus pneumoniae*^8,42^.

We analyzed the population structure of *C. diphtheriae sensu lato* in order to define phylogenetically informative LIN code thresholds. Our analysis showed a star-like phylogenetic structure, as most allelic differences corresponded to the upper and lower peaks in the distribution, separated by a wide plateau with few allelic distance values. This previously reported structuration into multiple deeply branching sublineages^3,5^, some of which have undergone recent clonal expansions, is thus confirmed here, based on a higher number of genomes and sublineages (from 114 sublineages represented previously, to 388 SLs herein). The creation and integration of a 10-level cgMLST-based LIN code system in the BIGSdb-Pasteur platform, enabled us to address various evolutionary and epidemiological questions.

Exploring deep evolutionary questions can be facilitated by using LIN codes, as illustrated here by the diversity of *tox* alleles among sublineages, which showed that only a few *tox* alleles were highly prevalent and showed broad distribution across sublineages. The *tox* gene distribution itself varied within some clonal groups, such as ClG8, but was fully compatible with the genetic cluster classification level, suggesting a good predictive power of this classification level for clinical implications.

The LIN code system also proved effective for epidemiological purposes at local and global levels. The escalating discrimination from genetic clusters (25 allelic mismatches) to complete LIN codes (0 allelic mismatches) is useful to reveal the finer-grained genetic diversity within emerging subpopulations resulting from recent clonal expansions. For example, studying the NTTB SL10037 protracted local transmission^4^, LIN codes revealed a large diversity with three clonal groups having diversified from a common ancestor, consistent with long-term local transmission from a reservoir of colonization and infections. On a broader scale, LIN codes illustrated that the epidemiological situation in New Caledonia and Yemen was characterized by diverse *C. diphtheriae* populations suggesting persistent endemic contexts (Figure 6, A and B). When investigating previously reported outbreaks, the analysis of LIN codes revealed distinct epidemiological dynamics, with more diversity having accumulated within SL8 (since the USSR 1990s outbreak) than within the recent expansions (Europe, 2022-2023) of SL574 and ST384 (Figures 6C, 6D and 6E). More diverse populations may indicate a longer diversification time, a more heterogeneous population influx, or a higher genetic mutation rate. In sum, the inspection of LIN codes within a given population of interest can uncover recent emergence or endemism, shifts in strain prevalence, and the spread of subtypes, which should be monitored in priority for their antibiotic resistance or virulence. Broader geographic application of genomic sequencing for diphtheria surveillance will help defining the relative roles of spatial spread and importation *versus* endemism of local strains, which currently appears as an important feature of *C. diphtheriae* populations.

**Figure 6.**
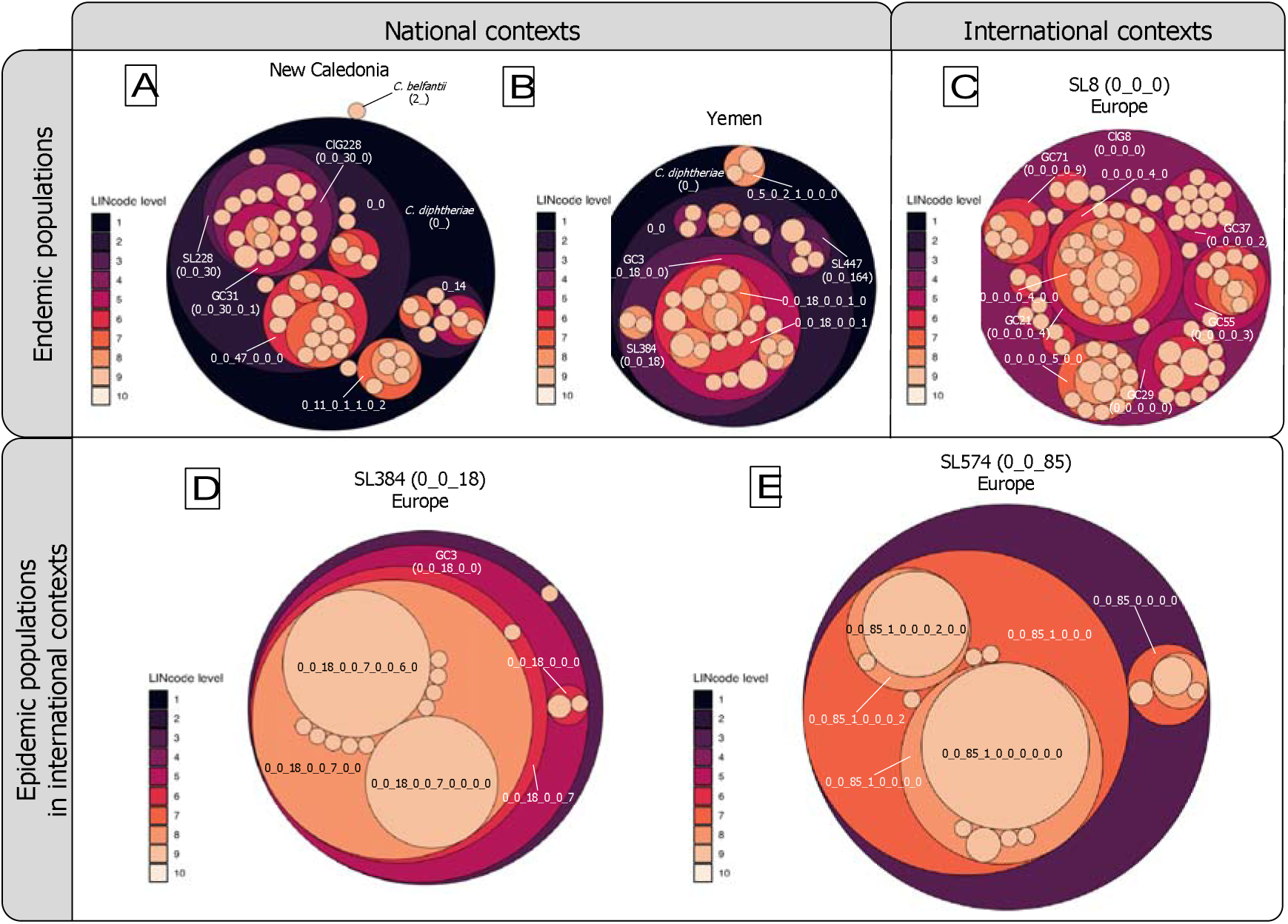
LIN code-based graphic representations of *C. diphtheriae* populations from different geographic and epidemiological contexts. LIN code prefixes are represented by circles coloured according to the level (from 1 to 10, as shown in the left legend of the panels), which are nested within their corresponding higher/parental LIN code prefixes. The circle sizes display the number of genomes encompassed by each LIN code prefix, and the labels for the main LIN code prefixes are displayed in the plots, including the nicknames for levels 1 and 3 to 5. The headings at the top and the left of the figure indicate the geographic and epidemiological contexts, and the title of each panel specifies the geographical region and/or the *C. diphtheriae* sublineage. **A:** Endemic *C. diphtheriae sensu lato* population in New Caledonia. **B:** Endemic *C. diphtheriae* population in Yemen (including GC3 population). **C:** Endemic (post-epidemic) SL8 population in Europe. **D:** Epidemic SL384 population in Europe (driven by epidemic GC3 population). **E:** Epidemic SL574 population in Europe (including epidemic GC1 population, which is covered by the fully overlapping 0_0_85_1_0_0_0 lower-level circle).

## Conclusions

We suggest that the adoption of the cgMLST-based LIN code system could mark a transformative step for the genomic taxonomy of *C. diphtheriae*. This system establishes a consistent framework for classification, while at the same time effectively mitigating the issues of instability caused by the merging of distinct groups associated with previously-used single-linkage clustering^3^. The LIN code system provides stability of taxonomic assignments, even at shallow levels, and enables differentiation among closely related isolates, which is essential to unify epidemiological tracking and outbreak investigations from different studies. Besides, the capability to elucidate relationships at different phylogenetic levels enriches our understanding of the evolutionary dynamics of *C. diphtheriae sensu lato*. The transition to the LIN code system should streamline communication on *C. diphtheriae* strains in research and public health contexts, ultimately contributing to more effective strategies for monitoring and managing this reemerging bacterial pathogen and the come-back of diphtheria.

## List of abbreviations

ANI: average nucleotide identity
BIGSdb: Bacterial Isolate Genome Sequence Database
CdSC: *Corynebacterium diphtheriae* species complex
cgMLST: core-genome Multi-Locus Sequence Typing
cgST: core-genome sequence type
ClG: Clonal group
DT: diphtheria toxin
GC: Genetic cluster
LIN code: Life Identification Number code
LIN tree: Life Identification Number-based tree
LMIC: low- and middle-income country
MALDI-TOF: Matrix Assisted Laser Desorption Ionization - Time of Flight
MCMC: Markov Chain Monte Carlo
MLST: Multi-Locus Sequence Typing
MRCA: most recent common ancestor
MST: minimum spanning tree
NTTB strain: non-toxigenic *tox*-bearing strain
PCR: polymerase chain reaction
SL: Sublineage
SNP: single nucleotide polymorphism
ST: sequence type

## Declarations

## Availability of data and materials

All genomes, allelic profiles and datasets analysed during the current study are available at the *Corynebacterium diphtheriae* species complex (CdSC) BIGSdb-Pasteur database (https://bigsdb.pasteur.fr/diphtheria/), including:

- Badell, E. *et al.* Ongoing diphtheria outbreak in Yemen: a cross-sectional and genomic epidemiology study. *Lancet Microbe* **2**, e386–e396 (2021): https://bigsdb.pasteur.fr/cgi-bin/bigsdb/bigsdb.pl?db=pubmlst_diphtheria_isolates&page=project&project_id=3
- Hennart, M. *et al.* A global *Corynebacterium diphtheriae* genomic framework sheds light on current diphtheria reemergence. *Peer Community J.* **3**, (2023): https://bigsdb.pasteur.fr/cgi-bin/bigsdb/bigsdb.pl?db=pubmlst_diphtheria_isolates&page=project&project_id=14
- Tessier, E. *et al.* Genomic Epidemiology of *Corynebacterium diphtheriae* in New Caledonia. *Microbiol. Spectr.* **11**, e0461622 (2023): https://bigsdb.pasteur.fr/cgi-bin/bigsdb/bigsdb.pl?db=pubmlst_diphtheria_isolates&page=project&project_id=10
- Hoefer, A. *et al.* Phenotypic and genomic analysis of a large-scale *Corynebacterium diphtheriae* outbreak among migrant populations in Europe. 2023.11.10.23297228 Preprint at https://doi.org/10.1101/2023.11.10.23297228 (2023): https://bigsdb.pasteur.fr/cgi-bin/bigsdb/bigsdb.pl?db=pubmlst_diphtheria_isolates&page=project&project_id=17

## Authors’ contributions

JFDB conducted the analyses and created the figures, with input from SB. MRP contributed to the implementation and curation of the LIN codes in the BIGSdb-Pasteur database. SB acquired funding and supervised the work. JDB and SB wrote the manuscript.

## Acknowledgements

We thank Virginie Passet for curating data in the BIGSdb-Pasteur diphtheria database, Federica Palma and Solène Cottis for BIGSdb-Pasteur project management assistance, and Brice Raffestin and Bryan Brancotte for maintenance and installing updates of BIGSdb-Pasteur. We thank Keith Jolley (Oxford University) for continuous developments of the BIGSdb platform, including LIN codes functionalities.

