## Additional File 1 for "Life identification number (LIN) codes for the genomic taxonomy of *Corynebacterium diphtheriae* strains"

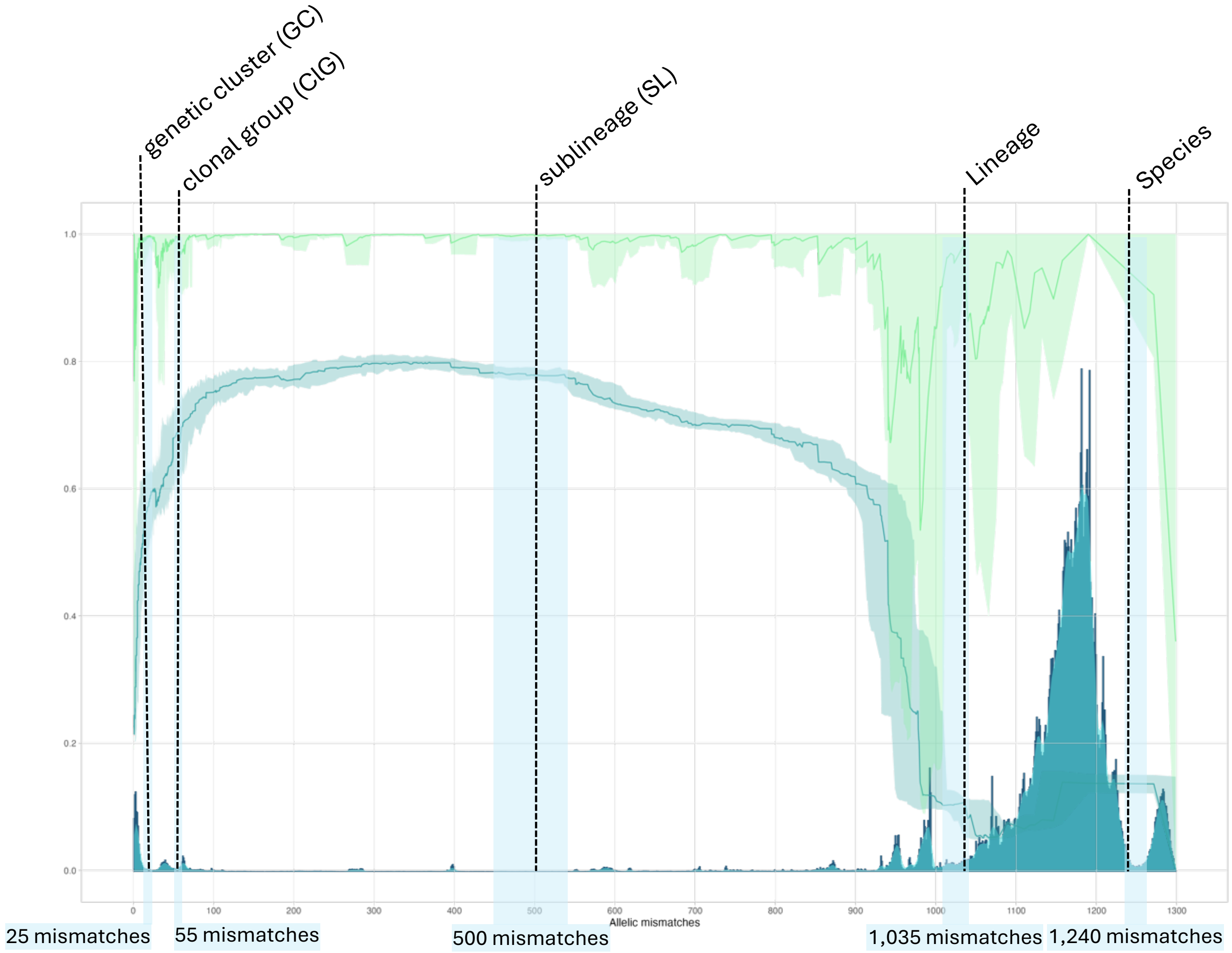


**Additional file 1.** Distribution of pairwise cgMLST distances, clustering properties, and phylogenetic congruence of *C. diphtheriae sensu lato* population. Threshold values ($t$) are shown on the $X$-axis, corresponding to allelic profile mismatch values up to 1,305 (or 100%). The dark blue histograms and the overlapping green curves display the distribution of pairwise allelic mismatches. The curves represent the consistency coefficient $S_t$ (silhouette, turquoise) and stability coefficient $W_t$ (Wallace, light green), respectively, obtained for each threshold $t$ (the corresponding scale is on the left $Y$-axis). To identify the two curves without reference to their colour, note that the $S_t$ curve starts (at $X$=0) approximately at 0.2 and the $W_t$ curve starts at 1.0. The light blue areas indicate the ranges of local optima: maximum $S_t$ values, minimum $W_t$ values, and low allelic mismatch counts. The dotted vertical black lines show the allelic mismatch thresholds (bottom labels) up to which the genome pairs belong to the same taxonomic level (top labels).
