## Additional File 2 for "Life identification number (LIN) codes for the genomic taxonomy of *Corynebacterium diphtheriae* strains"

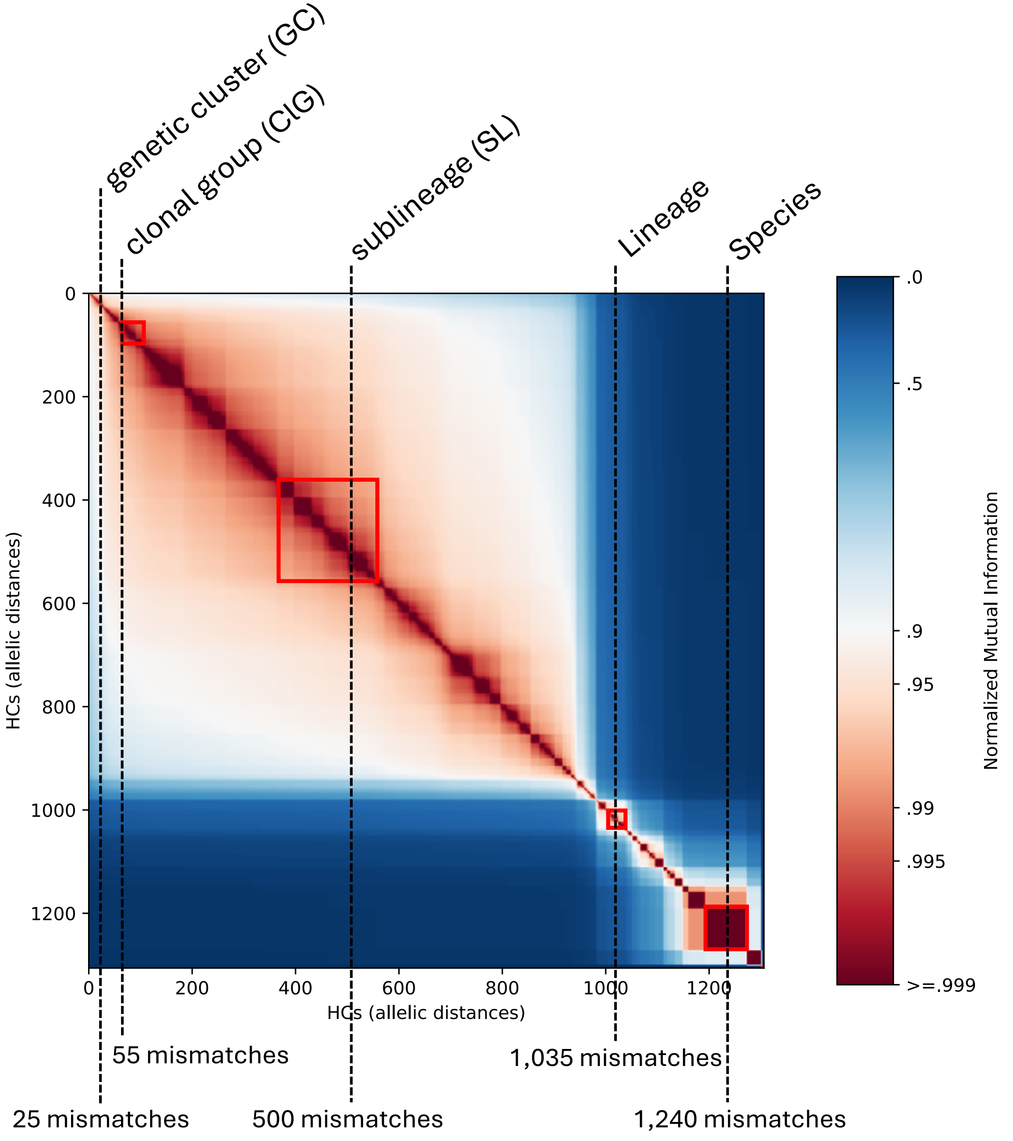


**Additional file 2.** Statistical evaluation of the cgMLST-based multi-level clustering from HierCC for *C. diphtheriae sensu lato*. The heatplot shows the Normalized Mutual Information (NMI) scores from pairwise comparisons of clusters at different hierarchical levels (from 0 to 1,305 pairwise allelic mismatches). The red boxes indicate the stable clustering ranges shared between HierCC and MSTclust, and the dotted black lines indicate the allelic thresholds (bottom labels) for *C. diphtheriae sensu lato* taxonomic levels definition (top labels).
