## Additional File 4 for "Life identification number (LIN) codes for the genomic taxonomy of *Corynebacterium diphtheriae* strains"

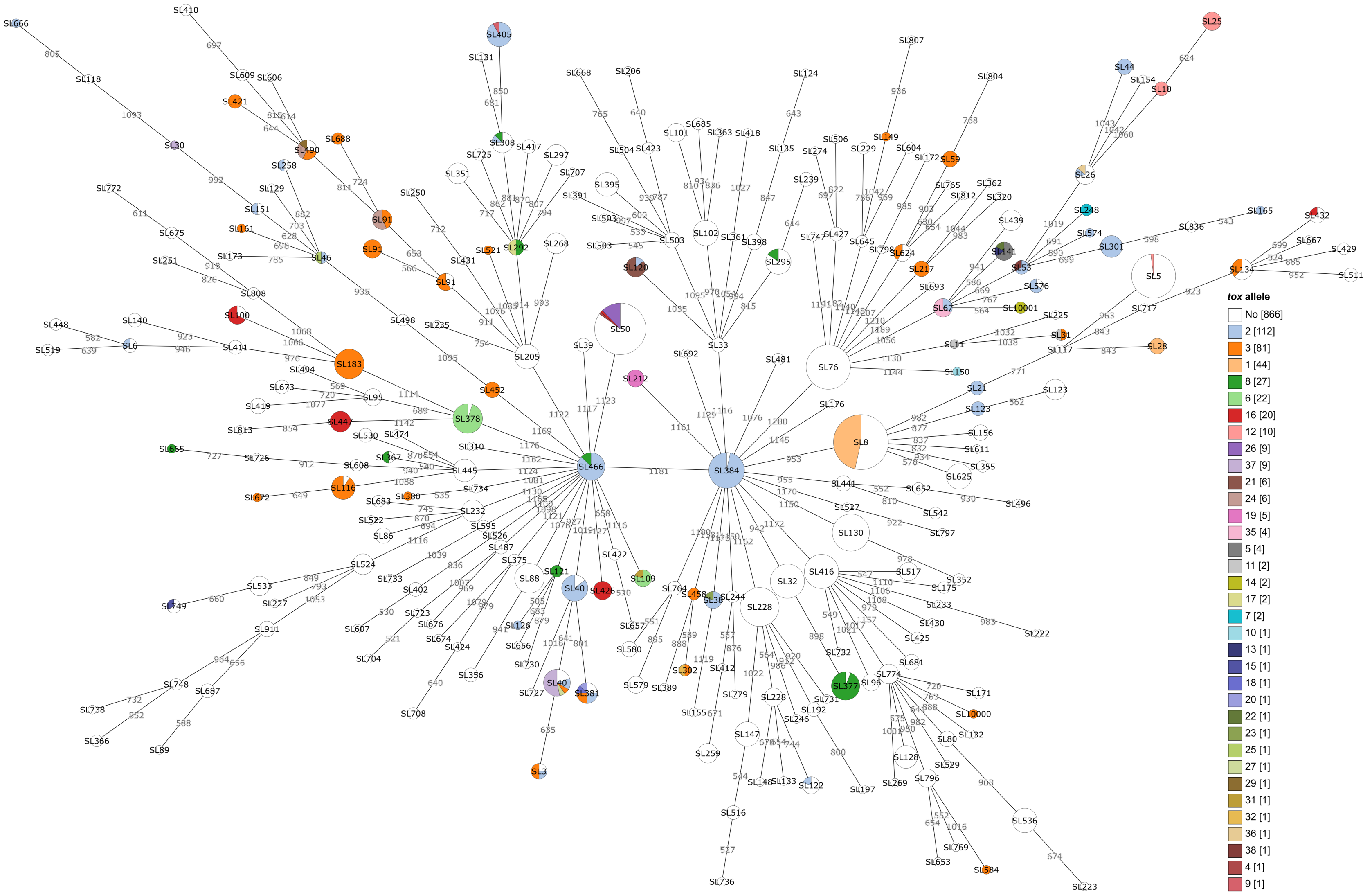


**Additional file 4.** Distribution of *tox* alleles in the *C. diphtheriae* population. Minimum-spanning tree generated with GrapeTree based on the *C. diphtheriae* cgMLST profiles of 1,250 genomes previously published[6]. The nodes represent different sublineages (SL, threshold of 500 allelic mismatches), which are indicated by labels on the nodes. The colours of the nodes indicate different *tox* alleles according to the right legend (*tox* allele [total number of allele-carrying genomes]). The sizes of the nodes are proportional to the number of genomes belonging to the indicated SL.
