## Additional File 5 for "Life identification number (LIN) codes for the genomic taxonomy of *Corynebacterium diphtheriae* strains"

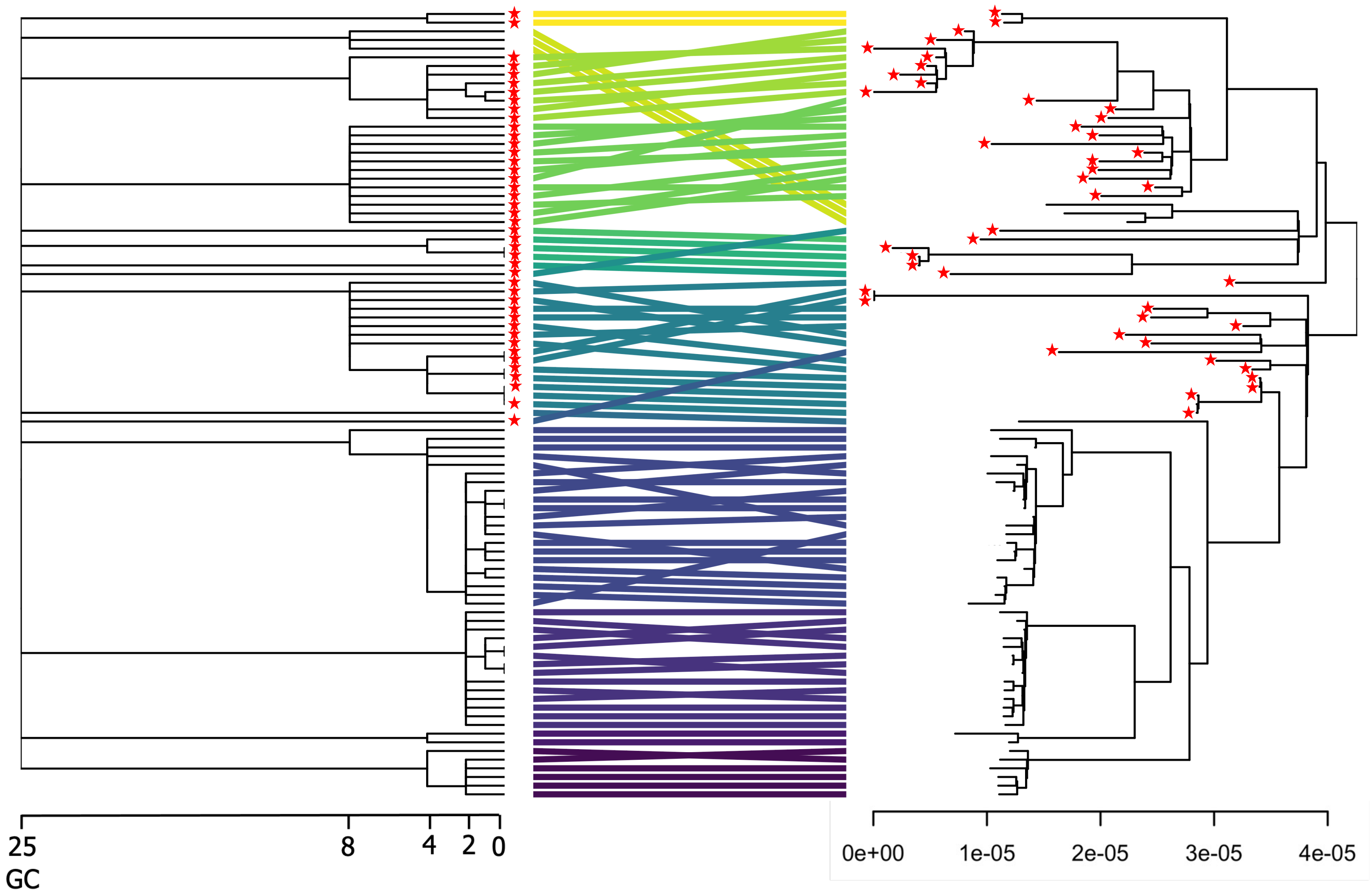


**Additional file 5.** Tanglegram showing the agreement between *C. diphtheriae* ClG8 trees. Left: LIN tree based on cgMLST LIN codes. The scale indicates the allelic mismatch thresholds from 25 (genetic cluster level) to 0 allelic differences. Right: Phylogenetic tree based on cgMLST SNPs. The scale indicates substituions per site. Links between tree tips are coloured according to the genetic clusters. Red stars in the tips of both trees indicate *tox*-positive isolates.
