## Additional File 6 for "Life identification number (LIN) codes for the genomic taxonomy of *Corynebacterium diphtheriae* strains"

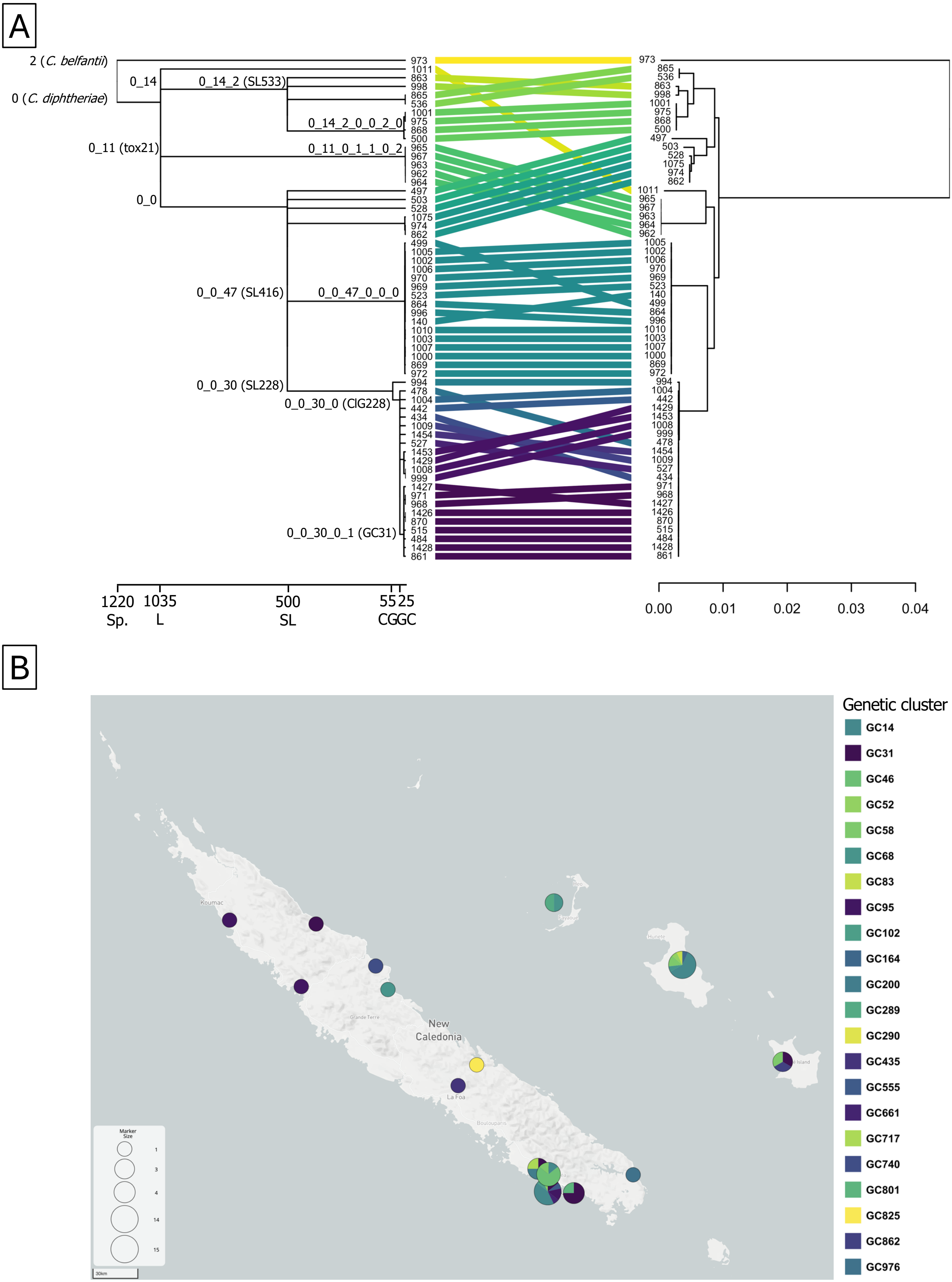


**Additional file 6.** LIN codes in the regional transmission of *C. diphtheriae sensu lato* from New Caldonia. **A.** Tanglegram showing the agreement between the LIN code-based LIN tree (left; scale indicates the allelic mismatch thresholds for the first five taxonomic levels) and the cgMLST SNP tree (right; scale indicates substituions per site) of a collection of 58 genomes from CdSC BIGSdb-Pasteur. Links between tree tips are coloured according to the genetic clusters (25 allelic mismatches). Nicknames and LIN code prefixes of the main species, lineages, sublineages (SLs), clonal groups (ClGs), and genetic clusters (GCs) are indicated next to the LIN tree branches. **B.** Geographical distribution of distinct genetic clusters in New Caledonia. The size of the nodes represents the total number of isolates from each sample location (left legend) and the coloured sectors represent the proportion of isolates belonging to each GC (right legend, same colours as panel A).
