## Additional File 7 for "Life identification number (LIN) codes for the genomic taxonomy of *Corynebacterium diphtheriae* strains"

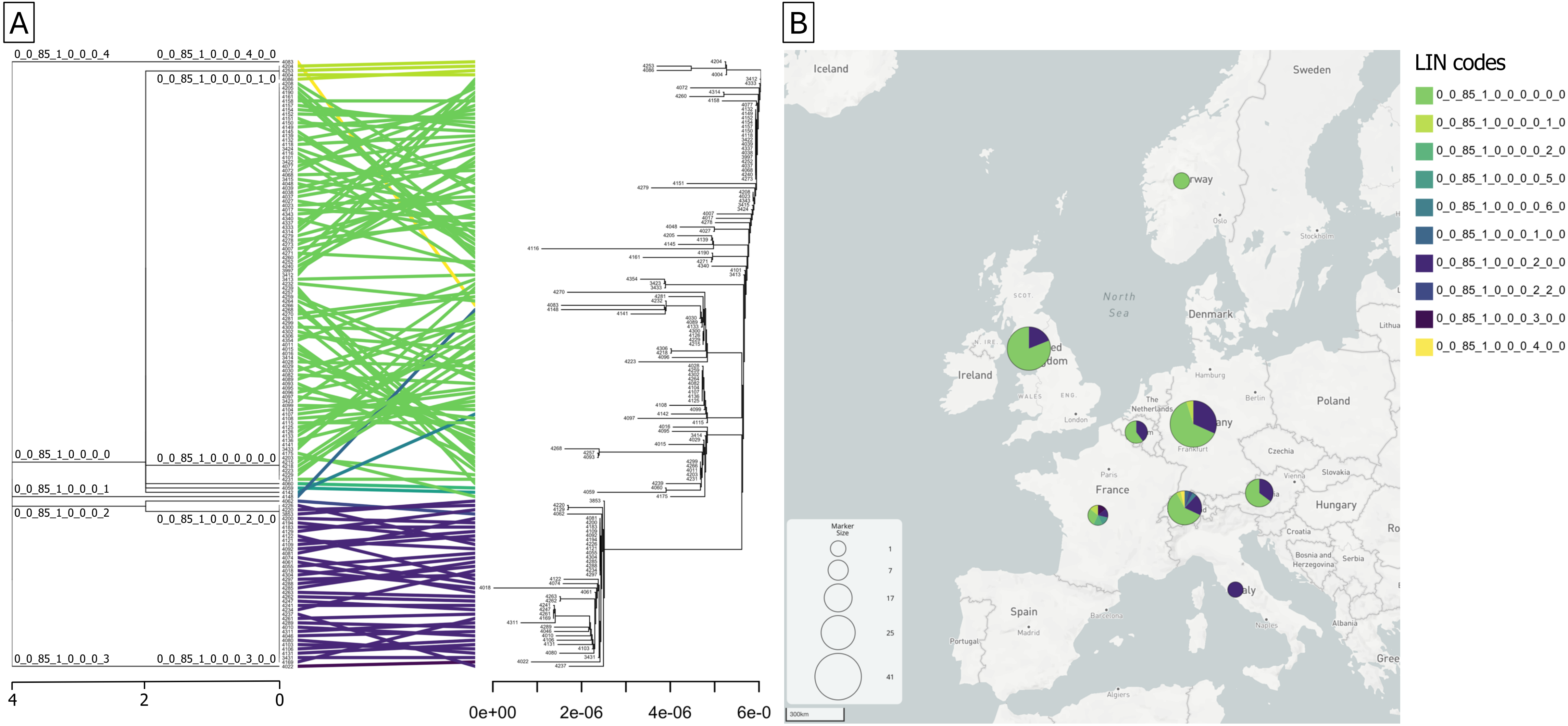


**Additional file 7.** LIN codes in the international ourbreak of *C. diphtheriae* GC1. **A.** Tanglegram showing the agreement between the LIN code-based LIN tree (left; scale indicates the allelic mismatch thresholds from 4 to 0 allelic differences) and the cgMLST SNP tree (right; scale indicates substituions per site) of a collection of 140 genomes from CdSC BIGSdb-Pasteur. Links between tree tips are coloured according to the complete 10-level LIN codes (0 allelic mismatches). LIN code prefixes of the main clusters are indicated next to the LIN tree branches. **B.** Geographical distribution of distinct GC1 clusters in Europe. The size of the nodes represents the total number of isolates from each country (left legend) and the coloured sectors represent the proportion of isolates belonging to each LIN code (right legend, same colours as panel A).
