## Additional File 8 for "Life identification number (LIN) codes for the genomic taxonomy of *Corynebacterium diphtheriae* strains"

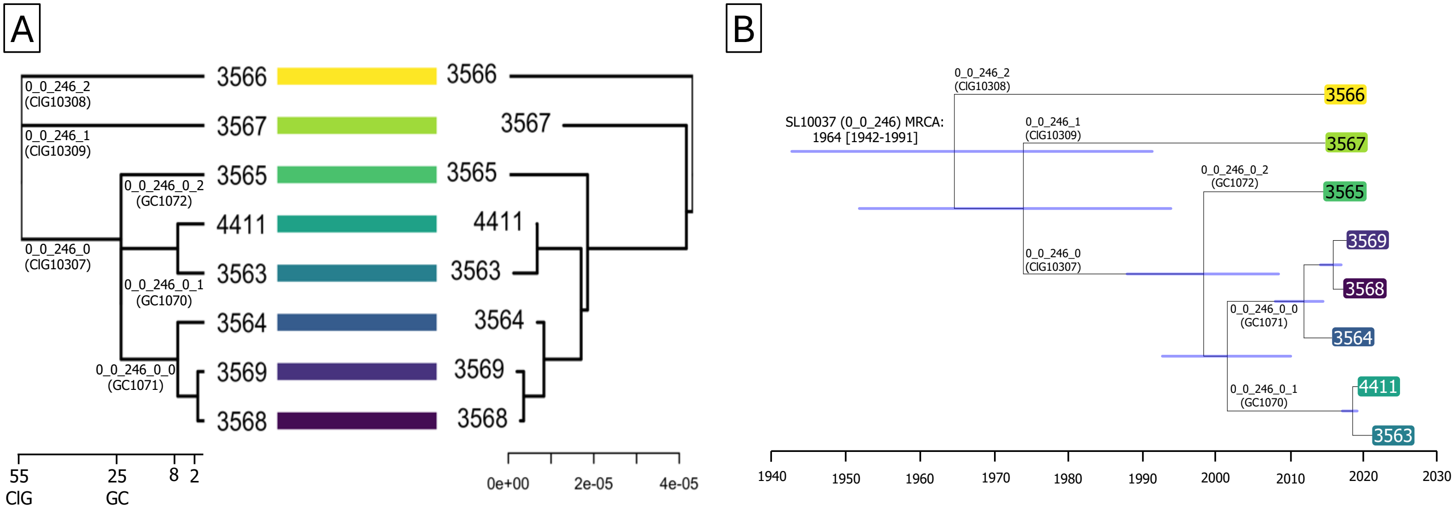


**Additional file 8.** LIN codes in the local transmission of NTTB *C. diphtheriae* SL10037 in England. **A.** Tanglegram showing the agreement between the LIN code-based LIN tree (left; scale indicates the allelic mismatch thresholds from 55 to 2 allelic mismatches) and the cgMLST SNP tree (right; scale indicates substitutions per site) of a collection of 8 genomes from BIGSdb-Pasteur. Links between tree tips are coloured according to complete 10-level LIN codes. Nicknames and LIN code prefixes of the main clonal groups (ClGs) and genetic clusters (GCs) are indicated next to the LIN tree branches. **B.** Dated phylogenetic tree of SL10037. SNP evolutionary rate: $1.3 \times 10^{-6}$ substitutions/site/year (95% CI $5.4 \times 10^{-7}$ to $2.4 \times 10^{-6}$). Confidence intervals (95%) for ancestral node dates are displayed by blue bars and indicated for the SL10037 most recent common ancestor (MRCA). Tip labels are coloured according to complete 10-level LIN codes. The main taxonomic clusters from sublineage to genetic cluster are indicated on the branches. The timescale is displayed in the *X* axis.
